# Enhancing Inter-link Coverage in Cross-Linking Mass Spectrometry through Context-Sensitive Subgrouping and Decoy Fusion

**DOI:** 10.1101/2023.07.19.549678

**Authors:** Boris Bogdanow, Cong Wang, Max Ruwolt, Julia Ruta, Lars Mühlberg, Wen-feng Zeng, Arne Elofsson, Fan Liu

## Abstract

In cross-linking mass spectrometry, sensitivity and specificity in assigning mass spectra to cross-links between different proteins (inter-links) remains challenging. Here, we report on limitations of commonly used concatenated target-decoy searches and propose a target-decoy competition strategy on a fused database as a solution. Further, we capitalize on context-divergent error rates by implementing a novel context-sensitive subgrouping strategy. This approach increases inter-link coverage by ∼ 30 - 75 % across XL-MS datasets, maintains low error rates, and preserves structural accuracy.

---

Cross-linking mass spectrometry (XL-MS) can reveal protein-protein interactions (PPIs) and the structural information of their binding interfaces in native biological systems ^1,2^. In XL-MS, protein contacts are captured using a cross-linker, a small organic molecule composed of a spacer arm and two reactive groups that typically target specific amino acid side chains, and are thereafter identified by mass spectrometry. Such experiments can yield intra-links (cross-links between residues within the same protein sequence), inter-links (cross-links between residues in different protein sequences) and mono-links (peptides that are modified by a partially hydrolyzed cross-linker). Inter-links only give insight into the structural configuration of PPIs.

The quality of XL-MS datasets critically depends on a balance between maximizing sensitivity (minimizing false negatives) and specificity (minimizing false positives). The same challenge is prevalent in standard bottom-up proteomics and is primarily addressed by estimating false-discovery rates (FDR) using a concatenated target-decoy search strategy^3^. In this approach, searches are performed against a database consisting of the target protein sequences and a concatenated list of reversed, shuffled, or randomized versions of them, called decoys ^3^. Any matches to the decoys are assumed to mimic false positives, which allows statistical control of error rates via the FDR. FDR filtering can be applied separately for different subsets of spectral matches when the error rates between the subgroups are reasonably different. Such or similar strategies are used for example for modified ^4,5^, miscleaved, or differently sized peptides in shotgun proteomics ^6^.

The XL-MS field has widely adopted and validated the concatenated target-decoy strategy^7–14^. Also, subgroup-specific FDR filtering is increasingly applied to separately analyze intra- and inter-links as inter-links have much higher error probability ^10,11^. However, this approach leads to low sensitivity in identifying inter-links. A potential solution to this challenge is considering that inter-links may have varying error probabilities dependent on the contextual information within the XL-MS dataset. In fact, recent studies show that selecting inter-links, where the corresponding proteins are additionally supported by intra-links or mono-links, significantly reduces the frequency of decoys on a concatenated database ^15,16^. Additionally, search engines were developed that integrate protein, PPI and cross-link level information into aggregate scoring functions to increase sensitivity ^13,14^. However, it has not been systematically assessed whether the basic assumptions of FDR control hold true when performing context-sensitive strategies.

Here, we find that currently applied concatenated target-decoy searches can drastically underestimate the FDR when considering the context of the dataset. Instead, we demonstrate that the theoretical error is better approximated by a fused decoy strategy. We leverage this observation to devise a novel subgroup FDR strategy that boosts inter-link identifications by 29 - 76 % in deep XL-MS datasets while simultaneously maintaining low error rates and increasing coverage of AlphaFold2-Multimer models across the proteome.

In order to investigate the suitability of a concatenated target-decoy strategy for context-sensitive strategies in XL-MS, we analyzed a deep XL-MS dataset from HEK293T cells, cross-linked with the enrichable cross-linker DSBSO ^17,18^ (“HEK”), yielding 12,216 target and 2,003 decoy inter-links when no cut-off is applied. Curiously, we observed 0 decoys among inter-links when both the inter-linked proteins contained an intra-link (i.e. context-rich subgroup) **(Figure 1a-b)**. To assess if this subgrouping strategy is valid, we simulated the distribution of true and false positives in this subgroup *in silico* based on a set of simple assumptions and controlled parameters **(see Methods)**. This is advantageous, as a simulation allows direct counting of the number of correct and incorrect matches **(Extended Data Figure 1a)**. In contrast to our initial observation, this simulation gives ∼1,300 false positives resulting in a fraction of false positives of 0.12 in this subgroup **(Figure 1b)**. This fraction is significant even when considering different database sizes and cross-link counts **(Extended Data Figure 1b)**. The problem becomes intuitively apparent when considering high numbers of intra-links in settings of restricted database size: When more and more intra-links are identified, many, if not most, proteins in a database will eventually contain an intra-link. Randomly assigned false positive inter-links will then more and more frequently assign proteins with these intra-links **(Figure 1c)**. We conclude that target-decoy competition with concatenated databases is unsuitable for controlling the FDR in XL-MS upon this type of subgrouping.

**Figure 1.**
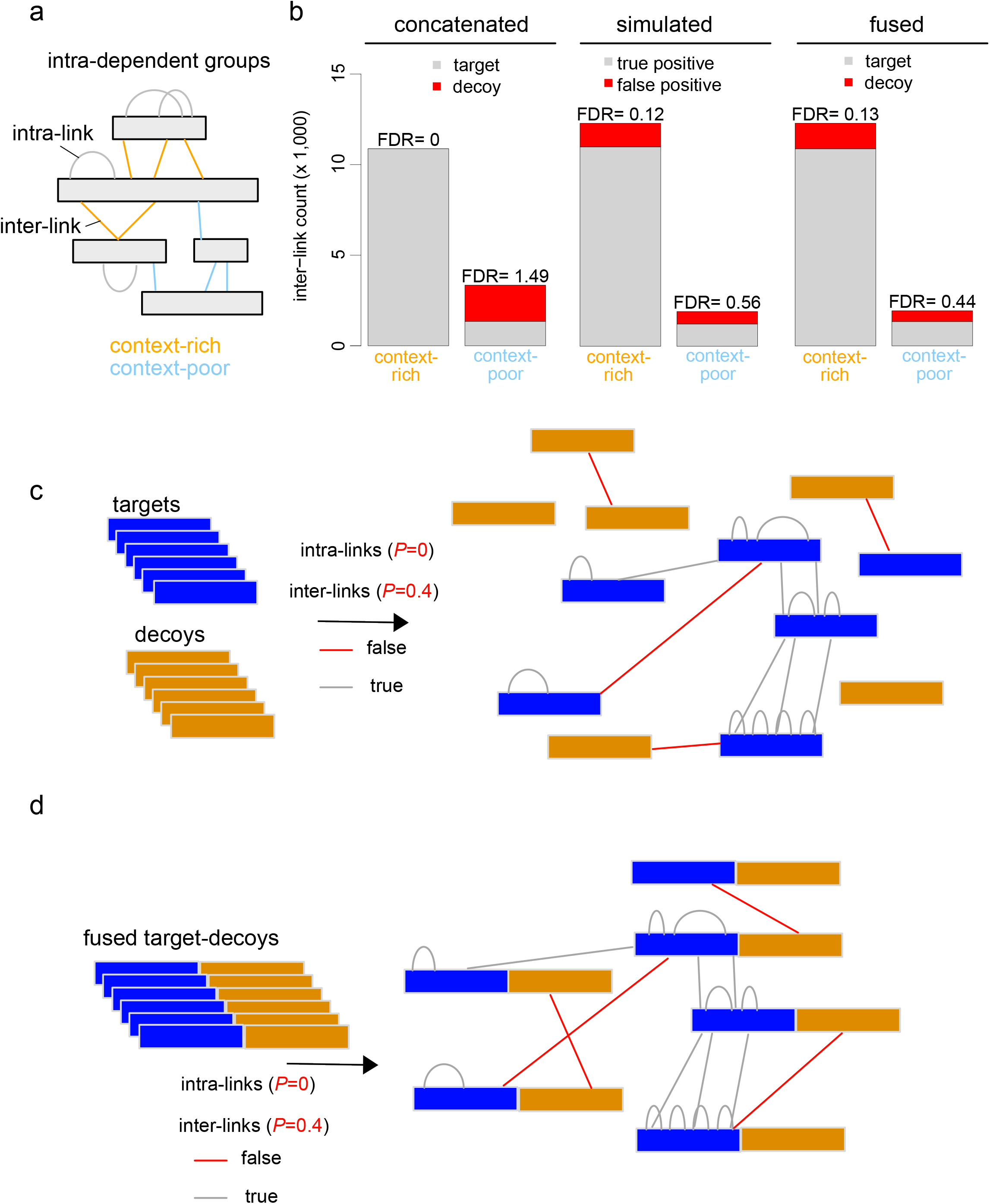
Accurate subgroup false discovery rates through decoy fusion. **(a)** Schematic of intra-dependent subgrouping. When both proteins were explained by intra-links, the corresponding inter-links were considered as a separate subgroup. **(b)** Subgroup false discovery rates upon concatenated or fused grouping strategies along with simulated fraction of true and false positives. FDR > 1 indicates higher decoy than target count (see calculation in the methods). **(c,d)** Schematic illustration of false and true positive assignments in concatenated **(d)** and fused **(e)** database search strategies. Overall intra- and inter-link count is 10 and probability for incorrect links is set to 0.4 and 0 for inter-and intra-links, respectively.

The problem arises as higher-level information (e.g. protein-level) is used to classify lower level matches (e.g. inter-links). Fused target-decoy databases, first employed in the shotgun proteomics analysis pipeline PEAKS DB^19^, can account for using higher-level information in the validation of lower-level matches. In this setup, each decoy entry is fused to its respective target entry, creating one fused protein sequence **(Figure 1d)**. In a standard, ungrouped setting, the distribution of target and decoy hits is not different when using a concatenated or fused decoy database. However, upon context-sensitive grouping, the usage of fused databases becomes essential as it ensures that once a target protein is assigned to a context-rich subgroup its corresponding decoy part is automatically assigned to the same subgroup. Repeating our previous analysis with a fused decoy strategy resulted in a higher and more realistic decoy count **(Figure 1b)**. The fused target-decoy estimates are also in line with the simulated data of two other datasets from purified mitochondria (“mito”)^20^ and herpesviral particles (“virion”)^21^ **(Extended Data Figure 1c)**, confirming that they adequately approximate the theoretically expected fraction of false positives.

Next, using the fused decoy database, we sought to further advance PPI coverage by introducing additional subgrouping strategies. To this end, we turned again to the “HEK” dataset and analyzed the score distributions of different subgroups. This analysis confirmed that the previously applied XL-MS subgrouping strategies, separating intra-from inter-links and focusing on inter-linked proteins that were additionally identified by intra-links, result in different score distributions (**Extended Data Figure 2a-b**). In addition, we found that inter-links had higher scores when other inter-links target different lysines of the same interacting proteins (“inter-dependent groups”) (**Figure 2a, Extended Data Figure 2c**), likely reflecting different probabilities of identifying wrong matches. Using a simulation approach similar to described above, we confirmed that inter-dependent subgrouping on a fused database accurately estimates overall error rates across the three datasets (**Extended Data Figure 3)**. We noticed that the context-rich subgroup exhibited particularly low error rates. This can be explained by the low probability of two different inter-links coincidentally matching to the same two protein sequences, which becomes exceedingly rare as the search space expands.

**Figure 2.**
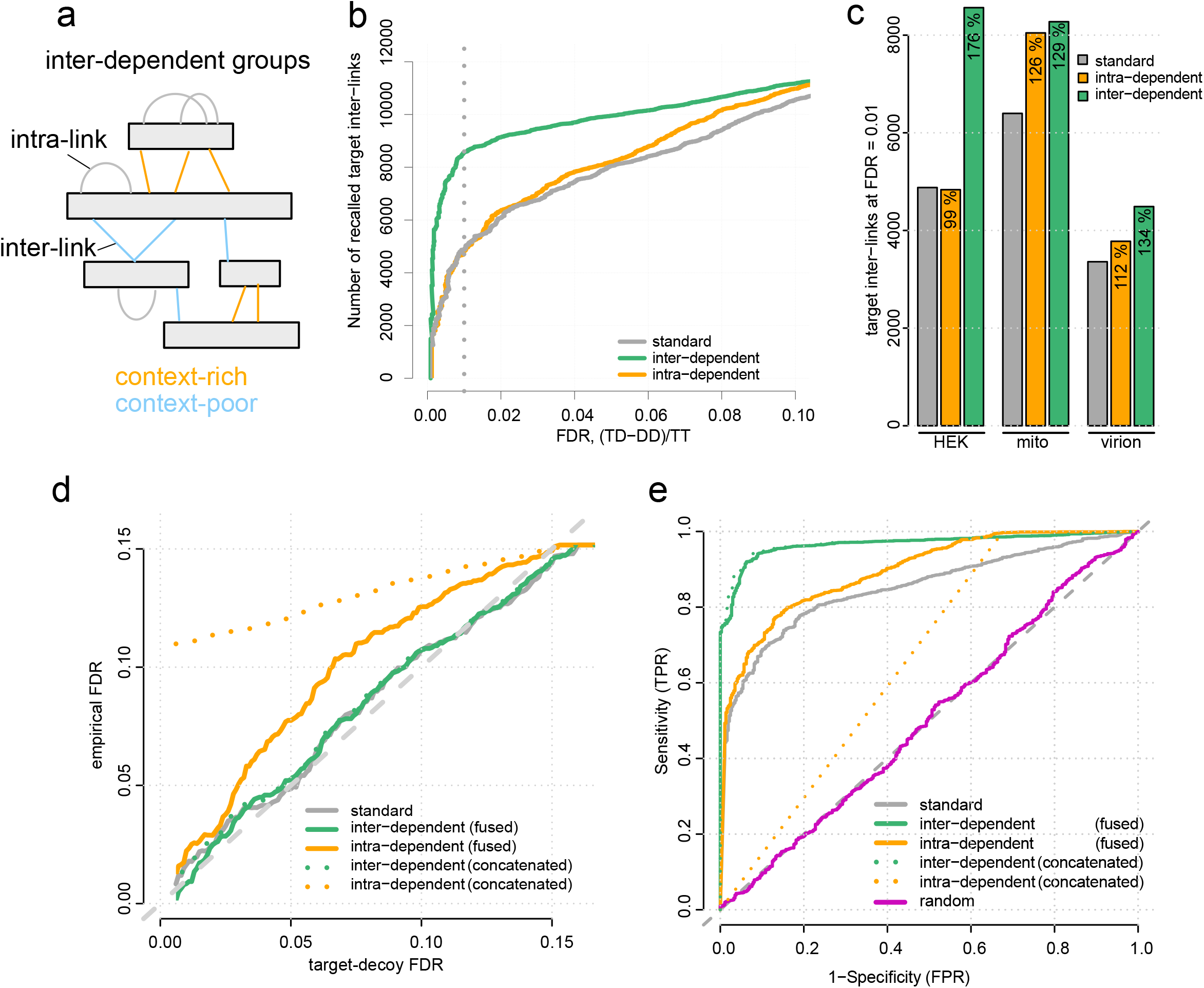
Improvement in statistical power through context-sensitive subgrouping. **(a)** Design of inter-dependent groups. When each protein in a PPI is targeted by at least two different lysines, the corresponding inter-links of the same PPI are considered a separate (context-rich) subgroup. **(b)** Recall of target inter-links as a function of the FDR cut-off comparing different strategies on a fused database for the HEK dataset. **(c)** Recall at 1 % FDR for three datasets and FDR strategies with relative change compared to the standard approach in percent. **(d)** Comparison of the target-decoy FDR to the empirical FDR as given by the mixing scheme in the ground-truth dataset for different FDR strategies. **(e)** Receiver operating characteristics (ROC) comparing different FDR strategies with varying decision thresholds to classify true and false positives.

To assess intra-dependent and the newly introduced inter-dependent subgrouping for their potential to increase sensitivity, we counted the number of identified inter-links that could be assigned at any FDR and compared these to the results of the standard search (that is, grouping all inter-links). In all tested datasets, we observed a pronounced increase in statistical power when inter-dependent grouping is applied **(Figure 2b, Extended Data Figure 4)**. The improvement was smaller (for “mito” and “virion” datasets) or non-existent (for “HEK”) upon intra-dependent grouping. In all cases, inter-dependent grouping performed best, particularly when error rates are low (29 - 76 % increase in sensitivity at FDR = 0.01) (**Figure 2c**). These data show that inter-dependent subgrouping increases sensitivity more robustly and effectively than intra-dependent subgrouping.

To further validate the benefits of the fused decoy strategy and inter-dependent subgrouping, we turned to our recently generated in vitro PPI dataset, which provides ground-truth empirical FDRs (Supplementary Note). As expected, the empirical FDR is well in agreement with the FDR based on target-decoy competition in the standard approach **(Figure 2d)**. Similarly, the proportion of false positives can be adequately controlled by inter-dependent grouping, regardless of whether concatenated or fused databases are used. However, when intra-dependent grouping is performed, only the fused decoy approach provides a reasonable (albeit consistently too low) estimate of the empirical FDR. Comparing all strategies using receiver operator characteristics (ROC) finds inter-dependent grouping as the best, with minor differences between fused and concatenated strategies (**Figure 2e**). We ascribe the excellent performance of this strategy to the fact that the inter-links belonging to PPIs that are additionally targeted by different inter-links are very unlikely to occur by chance. These very low error rates also prevent major differences between concatenated or fused approaches. Collectively, the results from three biological datasets, one controlled benchmark and our simulations prove context-sensitive subgrouping on a fused decoy database as a universally applicable strategy to increase sensitivity while maintaining accurate FDR estimates.

Finally, we assessed the utility of our optimized inter-link identification strategy for structural biology. In the “HEK” dataset, our strategy increased inter-link counts for 38 % of PPIs **(Extended Data Figure 5a)**. Mapping these onto proteome-wide predictions of complexes by AF2-multimer^22^, we find a similar fraction of model-satisfying inter-links compared to the standard approach **(Extended Data Figs 5b-d)**. The newly identified inter-links provide supporting structural information for both AF2-multimer models, such as the SMC2-SMC4 dimer (**Extended Data Figure 5e**, 10 additional cross-links) and large cryoEM structures, such as the proteasome **(Extended Data Figs 5f-g**, 17 additional cross-links).

Our study demonstrates that context-sensitive subgrouping in combination with a fused decoy search enhances the statistical power of XL-MS analyses and hence the coverage of PPI contact sites. Due to the inherent structure of XL-MS data, we see great potential of emerging context-sensitive strategies ^13–15^. Future work may be directed towards exploring other grouping criteria or aggregate scoring functions that integrate multiple levels of information to continue pushing the boundaries of system-wide structural PPI profiling.

## METHODS

### Generation of HEK dataset

We generated a large-scale XL-MS dataset from intact HEK293T cells using Azide-a-DSBSO (Sigma-Aldrich) as a crosslinker. HEK293T cells were grown to 90 % confluency on 15 cm cell culture dishes and harvested using ice cold PBS. Cells were washed twice with ice cold PBS before protein concentration estimation using ROTI®Quant. Cells were adjusted to a concentration of 10 mg/ml in PBS and crosslinked in vivo at room temperature using 2 mM Azide-a-DSBSO for 30 min. Quenching was performed with 30 mM TrisHCl pH 7.4 for 10 min. The cross-linked HEK293T cells were pelleted using a desk centrifuge (RT, 10 min, 5000 g). The cell pellets were solubilized using 8 M Urea in 50 mM TEAB pH 8.5 at room temperature and subjected to supersonic treatment using a Bioruptor (5 min, cycle 30/30 sec). 1 mM DTT and 0.5 ug benzonase per 10 mg cells were added to the lysate and the sample was incubated for 45 min at 37 °C under shaking at 1000 rpm. Afterwards, the reaction was quenched using 40 mM Chloroacetamide and the sample was incubated for 30 min at room temperature in darkness. Endopeptidase Lys-C was added to the sample 150 : 1 (protein : enzyme, w/w) and the sample was incubated 3 h at 37 °C under shaking at 1000 rpm. The Urea concentration was diluted to 2 M using 50 mM TEAB pH 8.5 and tryptic digest was performed overnight (100 : 1; protein : enzyme; w/w).

The digest was desalted using Sep-Pak columns and solubilized in PBS. The isolation of Azide-a-DSBSO crosslinked peptides was performed using DBCO agarose beads (Click-chemistry tools) overnight at RT. Afterwards, DBCO agarose beads were successively washed with SDS, 8M Urea in 50 mM TEAB pH 8.5, 10 % ACN, and ultrapure water. The crosslinked peptides were eluted for 2 h using 10 % TFA in ultrapure water. Crosslinked peptides were separated from monolinks using size exclusion chromatography on a Superdex 30 Increase column with 3.2 x 300 mm bead dimensions (cytiva) and the isolated crosslinks were subjected to a high pH HPLC run using a Gemini column 3µm C19, 110 LJ, 100 x 1mm (phenomenex) on an agilent offline fractionation system. The fractions from the high pH HPLC run were subjected to LC-MS analysis individually.

### LC-MS/MS

For the HEK dataset, LC-MS/MS analysis was performed using a Orbitrap LUMOS system from Thermo fisher scientific with an Ultimate 3000 HPLC system. Reversed-phase separation was performed using a 50 cm analytical column (in-house packed with Poroshell 120 EC-C18, 2.7 µm, Agilent Technologies) with a 180 min gradient. Spectra were recorded using FAIMS CV combination of (−50/−60/−70) and HCD-MS2 acquisition was performed using the normalized collision energies: 19, 25, 30 with a charge filter of +4 to +8. MS1 and MS2 scans were acquired in the orbitrap with a respective mass resolution of 120,000 and 60,000. Dynamic exclusion was set to 60 s and the isolation window was set to 1.6 m/z with a normalized AGC target of 200 %.

### Cross-link searches

Peak lists (.mgf files) were generated in Proteome Discoverer to convert each .raw file into one .mgf file containing HCD-MS2 data. The .mgf files were used as input to identify cross-linked peptides with a stand-alone search engine based on XlinkX v2.0 ^2^. The following settings of XlinkX were used: MS ion mass tolerance, 10 parts per million (ppm); MS2 ion mass tolerance, 20 ppm; fixed modification, Cys carbamidomethylation; variable modification, Met oxidation; enzymatic digestion, trypsin; and allowed the number of missed cleavages, 3; DSBSO cross-linker, 308.0388276 Da (short arm, 54.01056 Da, long arm, 236.01770 Da).

All MS2 spectra were searched against concatenated target-decoy databases (randomized decoys) generated based on the corresponding proteome determined by bottom-up proteomics, containing 4,860 target sequence entries. The table giving targets and decoy residue-level cross-links at no FDR threshold was used for subgrouping, FDR calculations and input for simulations.

### Target-decoy fusion strategy

In order to evaluate the performance of a fused decoy database search strategy, we analyzed our datasets as if they were searched using a fused database. To this end, we performed the database search in a concatenated design but then replaced the entries containing the gene names for the cross-linked proteins with their target annotations in the cross-link results table. For example, a cross-link matching the proteins “RPL12-RPS6(decoy)” was re-annotated as “RPL12-RPS6”. Thus, all targets and decoys originating from the same fasta entry are fused into one entry. Importantly, we kept an additional identifier indicating whether the cross-link originated from the decoy or target part of the fused entry.

### Subgrouping and FDR calculations

We devised only two subgroups for intra- and inter-dependent filtering to assure subgroups are sufficiently large to enable target-decoy competition^5^ and to avoid overfitting to a specific dataset. In the case of intra-dependent subgrouping we devised two subgroups for each strategy: One context-rich group containing all inter-links where both the cross-linked proteins are additionally supported by intra-links and a second, context-poor, subgroup containing inter-links where at most one of the proteins contained intra-links. In the case of inter-dependent grouping, we also devised two subgroups. A context-rich subgroup containing the inter-links where each of the proteins in a PPI is supported by at least two independent inter-linked lysines between the same proteins and a second, context-poor, subgroup containing all other inter-link matches. Groupings were performed at the level of residue-residue connections, and all residue-residue identifications without FDR control were used for subgrouping. Then the individual lists were sorted by decreasing search scores. We here used the lower (worse) negative decadic logarithmic XlinkX search scores reported for both sides of the cross-link as the overall score for the cross-link. The subgroup false discovery rate for each entry in the list in both the subgroups independently was then calculated based on Eq (1), where T denotes a target match, and D is a decoy match on either side of a cross-link, respectively.

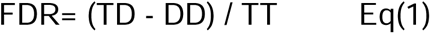

Then, subgroup false discovery rates from both groups were assembled into a combined list and sorted by their subgroup false discovery rates. We then performed another round of FDR calculations on each of the entries in the combined list according to Eq(1), giving the overall FDR. Standard (ungrouped) FDR calculations were performed based on Eq(1) on the total list of inter-link identifications based on the database search score (see above).

### Simulation

Two simulations were performed to estimate the proportion of false positives between subgroups. The estimation of false positives for intra-dependent grouping involved considering the following input parameters: (i) the database size *n* (the number of target entries used for search), the number of inter-links, the number of intra-links, and the probability of incorrect matches for inter-links (fraction decoy inter-links). These parameters were calculated based on a search table containing all cross-links (residue-residue connections) without any FDR control. To determine protein-dependent probabilities for wrong matches, a Poisson distribution was fitted to the observed decoy cross-link counts per protein. The resulting λ coefficient was utilized to generate a normal distribution with *n* counts, λ standard deviation, and λ mean. Protein-dependent match probabilities were calculated by dividing each value of the resulting distribution by the sum of all values. Values below 0 were set to 0. This simulation assumed that wrong matches occurred randomly, without considering the possibility of strong systematic biases towards specific proteins, such as through misidentified modified peptides ^23^. To estimate protein-dependent probabilities for correct matches, a Zipf distribution ^24^ was fitted to the observed target cross-link counts per protein. The resulting coefficient *s* was used to calculate the protein-dependent matching probability for each unique value under the fitted distribution. By utilizing a power-law distribution to model the correct cross-links, the nature of protein interactions in real-world datasets was captured. Power-law distributions account for the presence of a “long tail,” where a small number of highly connected proteins have a disproportionate impact on the network’s connections. Subsequently, incorrect and correct inter-links and intra-links were deposited onto the *n* entries according to these probabilities, and the fraction of incorrect matches was evaluated in the subgroups of inter-links where (i) both linked proteins contained at least one intra-link and (ii) all others.

For simulating the frequency of false positive matches in inter-dependent subgrouping, the simulation was adapted as follows: (i) intra-links were not considered, and (ii) inter-link matches were split into two parts, reflecting the possibility that each of the two cross-linked peptides could be wrongly matched (target-target, decoy-target, decoy-decoy). The frequency of these matches was obtained from their frequency in the actual datasets without FDR control. A Poisson distribution was fitted to the empirically observed counts of decoy matches for all cross-links per PPI involving the most frequently linked decoy protein. To simulate correct matches, a Zipf distribution was fitted for all cross-links per PPI involving the most frequently linked target protein. Incorrect and correct matches were then deposited on both sides of the inter-link according to the resulting probabilities onto the n entries (as described in the previous paragraph). The frequency of incorrect matches was evaluated in the subgroups of inter-links where (i) both of the linked proteins contained at least one additional inter-link between the same proteins and (ii) all others.

### AF2 prediction and structure mapping

Hetero-dimers were predicted by AlphaFold2 multimer v2.1^25^ using the default protocol and sequence search methods. For each model, the pDockQ score was calculated ^26^. Different model qualities were considered based on the pDockQ score. Mapping of cross-links on AlphaFold2-Multimer predictions was performed using the bio3d R package. Therefore, we extracted the C-alpha atom coordinates of inter-linked Lysines in the predicted zero rank dimeric structure and calculated their distance in three-dimensional space.

## DATA AVAILABILITY

The mitochondrial (DSSO) dataset is pre-printed ^20^ and available under PXD032132 (, password: XXXXX), The virion (DSSO) dataset is in press ^21^ and available under PXD031911. The HEK dataset has been deposited to the ProteomeXchange Consortium via the PRIDE ^27^ partner repository with the dataset identifier PXD043531 (, password: XXXXXX).

## CODE AVAILABILITY

Original code related to FDR calculations and simulations is provided under https://github.com/Bogdanob/XLMS_decoyFusion/.

## ACKNOWLEDGEMENTS

BB acknowledges funding from DFG grant BO 5917/1-1. CW, MR, JR are supported by the European Research Council (ERC) Starting Grant (ERC-STG No. 949184). LM is funded by the Leibniz-Wettbewerb (K284/2019). The AF2 computations were enabled by resources provided by the Swedish National Infrastructure for Computing (SNIC) at NSC (Berzelius), partially funded by the Swedish Research Council through grant agreement no. Berzelius-2021-29 and SNIC 2022/5-282 (AE).

## AUTHOR CONTRIBUTIONS

BB and FL initiated and designed the project, and conceptualized the FDR filtering strategies. BB performed most analyses and conceptualized decoy fusion and inter-dependent grouping for XL-MS. CW prepared the HEK dataset. MxR contributed to analyses of the ground-truth dataset. WZ contributed to FDR strategy discussions. JR and LM contributed to structure mapping. AE predicted AF2 multimer models. BB and FL wrote the manuscript, AE made minor suggestions to the manuscript.

---

**Extended Data Figure 1.**
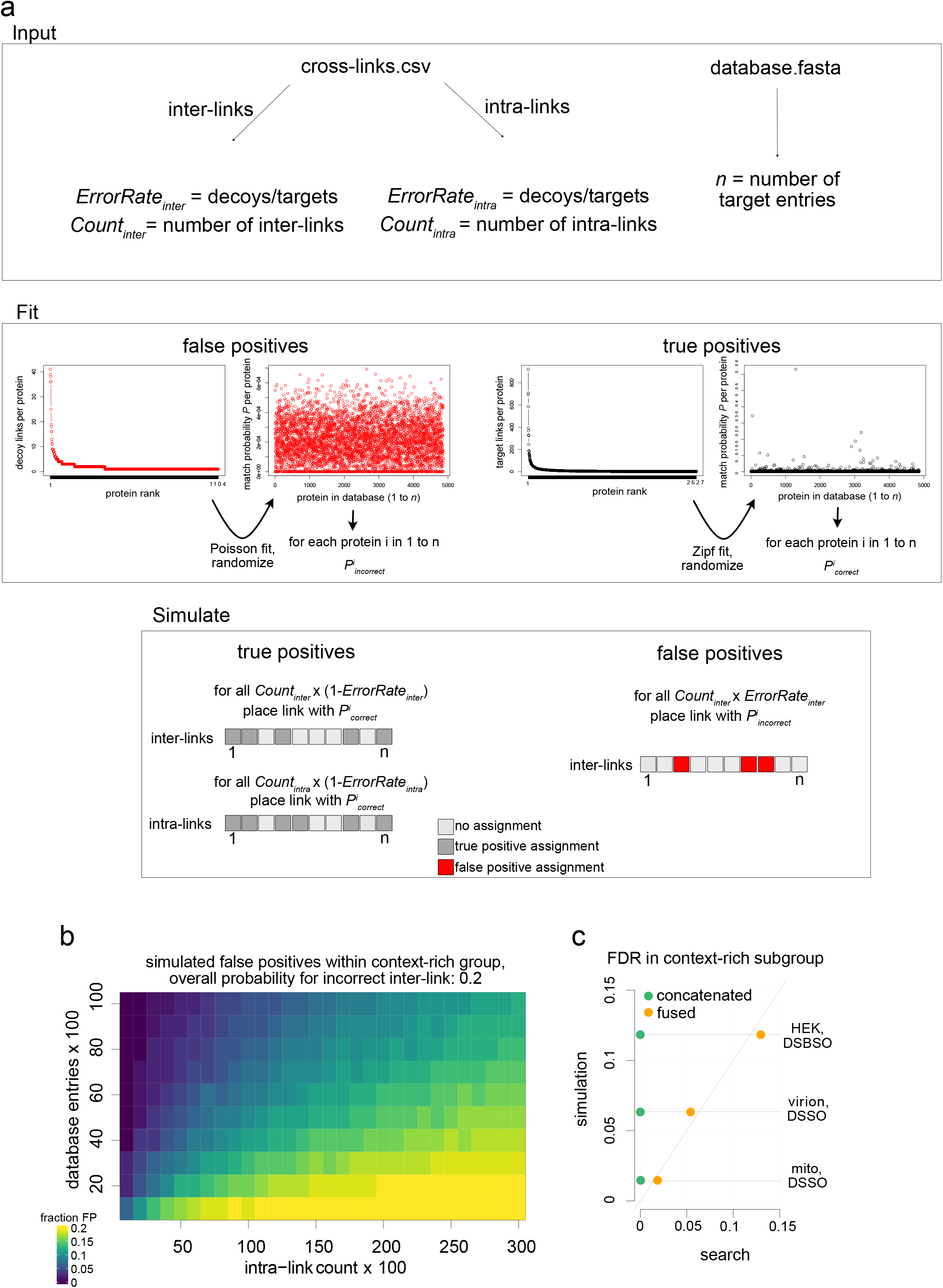
Simulating false positives in XL-MS datasets. **(a)** Workflow for simulations. Database size, the number of cross-links (inter and intra) and their decoy/target fractions were used as input. Protein-dependent probabilities for wrong matches were obtained from a Poisson fit through the decoy cross-link count per protein. Protein-dependent probabilities for correct matches were obtained from a Zipf fit through the target cross-link count per protein. Correct and Incorrect cross-links were deposited according to these probabilities on the proteins in the database. **(c)** Simulated fraction of false positives in the context-rich subgroup (both proteins have intra-links) in dependence of database size and intra-link count. Overall probability for obtaining an incorrect inter-link set to 0.2. In cases of restricted database size and high intra-link count the overall (ungrouped) error rate is approached. **(d)** Inter-link FDR in the context-rich subgroup upon simulation, and upon fused or concatenated search for three biological datasets.

**Extended Data Figure 2.**
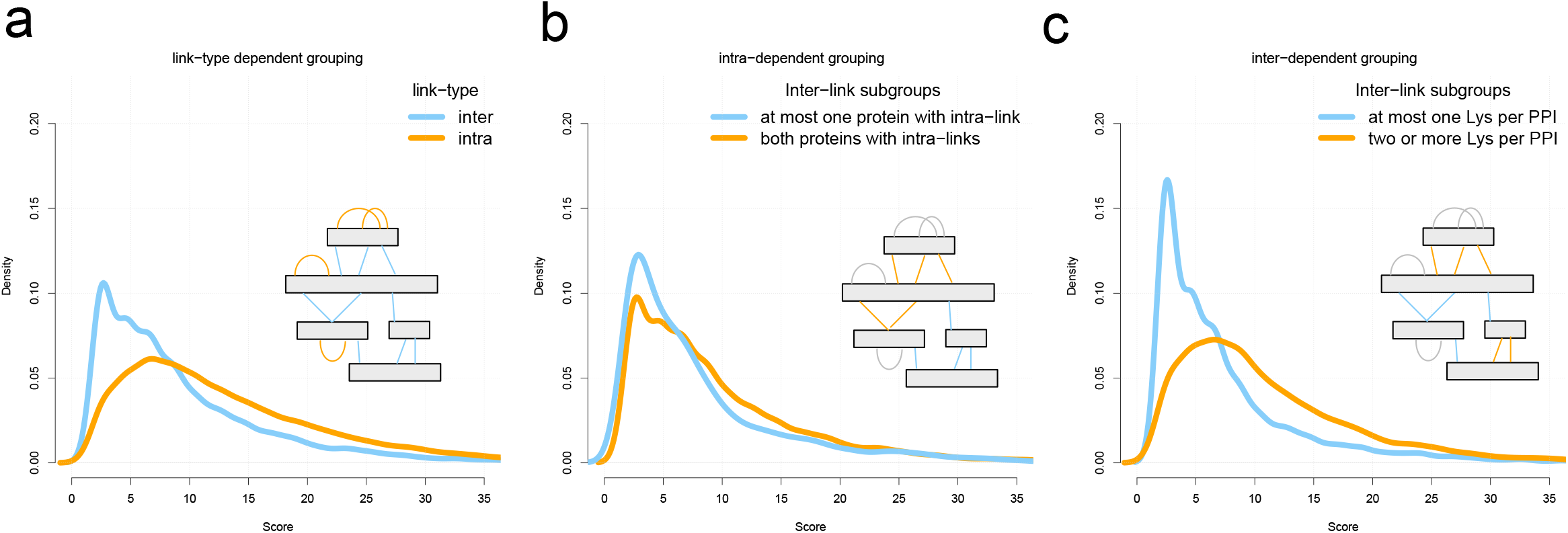
Cross-link score distributions in different subgrouping strategies. Cross-link search scores (negative decadic logarithmic XlinkX score) upon **(a)** standard subgrouping separating intra-from inter-links **(b)** inter-dependent subgrouping and **(c)** intra-dependent subgrouping in HEK dataset, all grouped on a fused database.

**Extended Data Figure 3.**
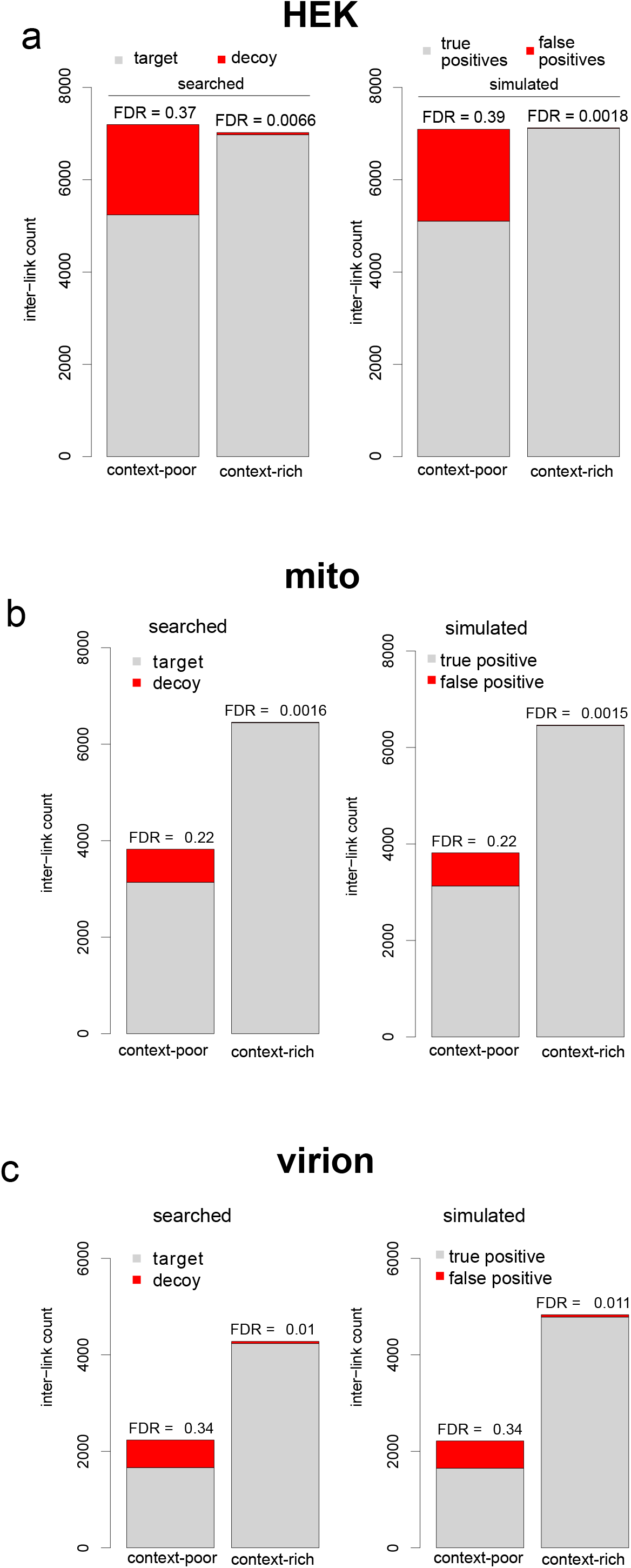
Inter-dependent grouping agrees with simulated error rates. Inter-link target and decoy count in inter-dependent subgroups and fused strategy for **(a)** HEK **(b)** mito and **(c)** virion dataset along with inter-link true and false positives obtained from simulated distributions.

**Extended Data Figure 4.**
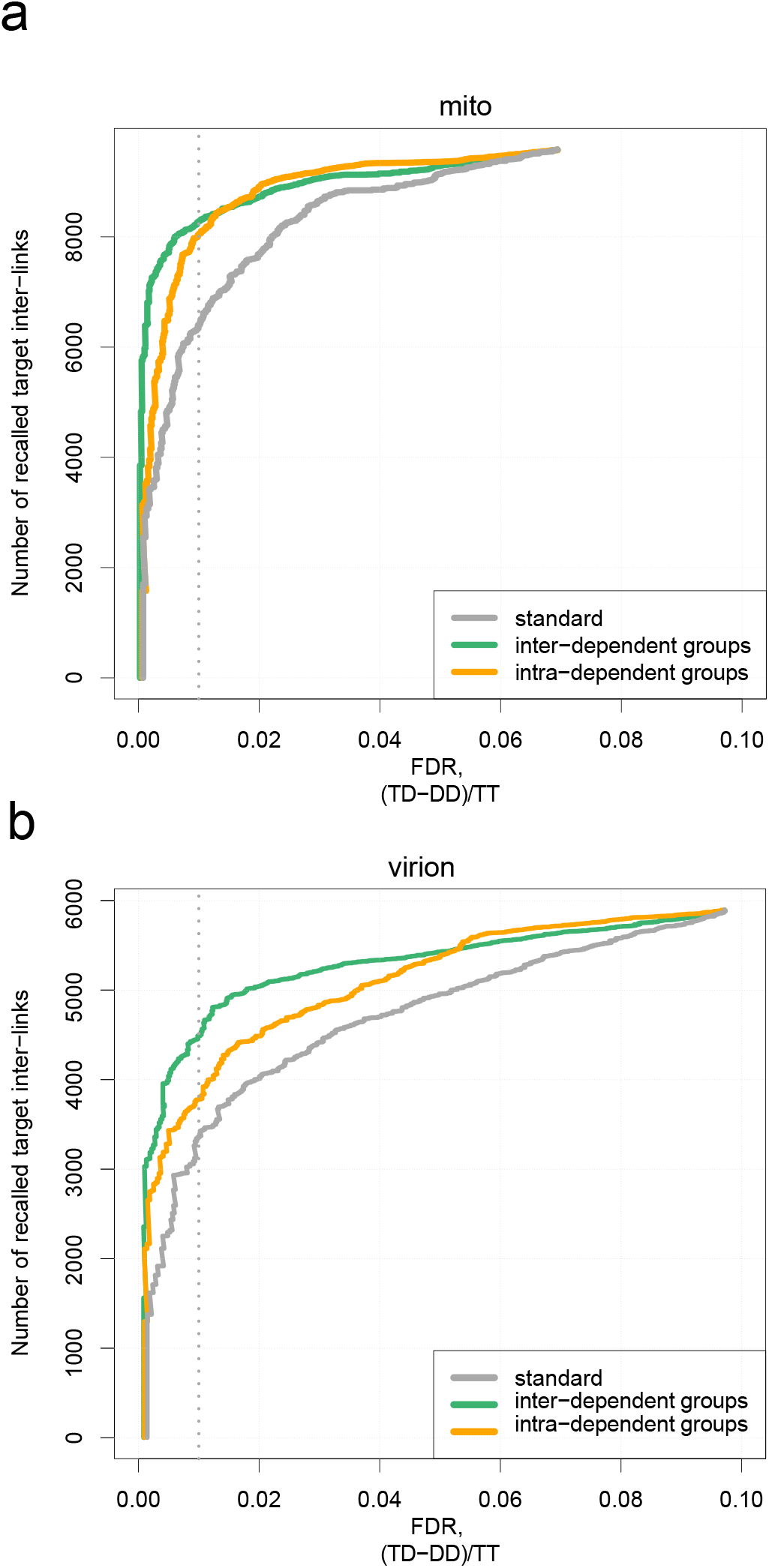
Improvement in statistical power through context-sensitive subgrouping. Recall of target inter-links as a function of the FDR cut-off comparing different strategies on a fused database for the **(a)** mito and **(b)** virion datasets.

**Extended Data Figure 5.**
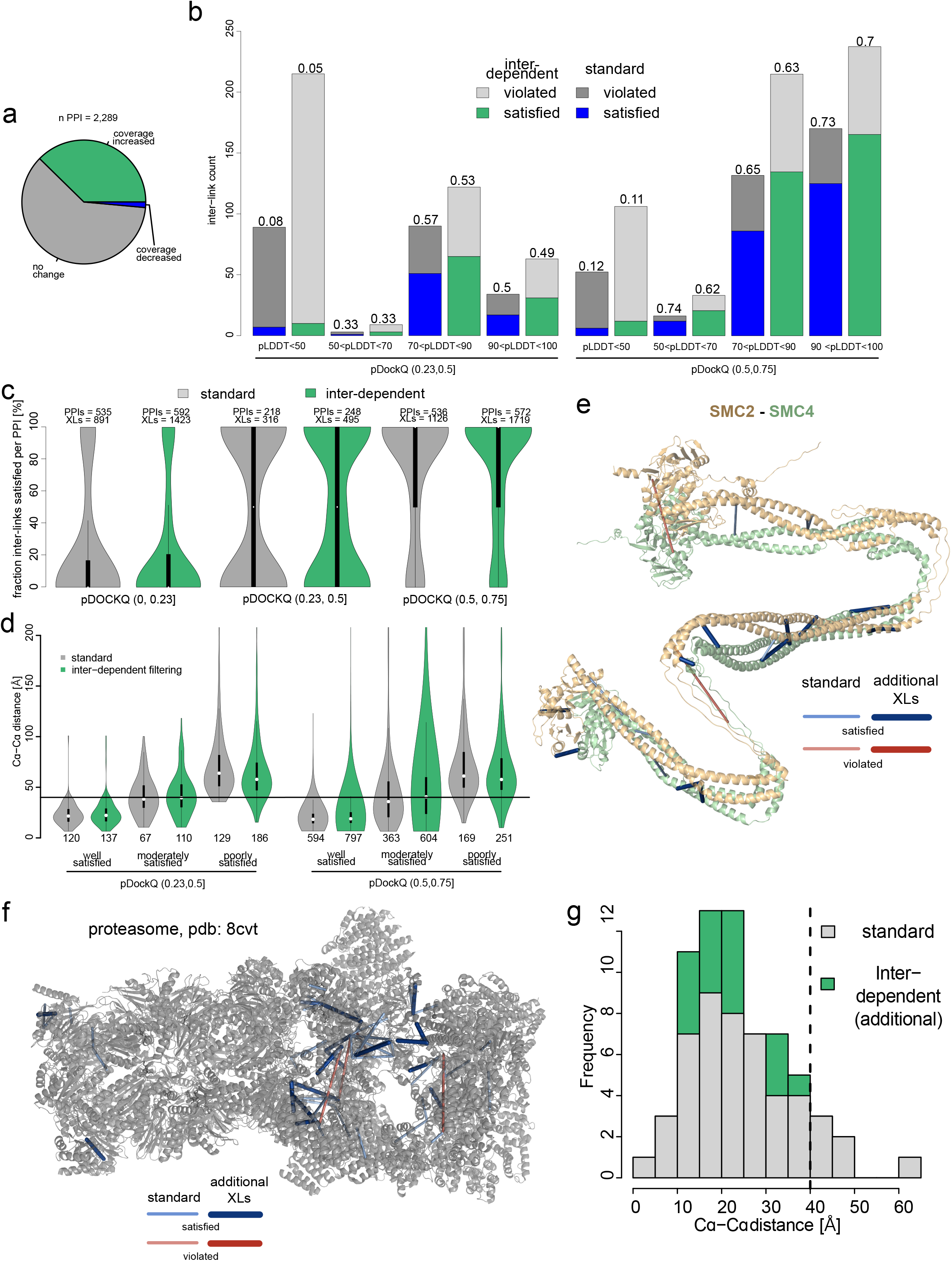
Structural accuracy of inter-dependent subgrouping. **(a)** Differences in inter-link coverage of PPIs due to inter-dependent subgrouping compared to standard. **(b)** Number of Inter-links violating and satisfying the distance constraint of 40 □ comparing inter-dependent subgrouping and standard approach (grouping all inter-links). Varying thresholds of model quality (pDockQ) and residues (pLDDT, on both Lysines) were considered. **(c)** Fraction of inter-links per PPI satisfying the distance constraint comparing both strategies for poor, moderate and well-predicted models. A pLDDT cut-off of 50 was applied for both cross-linked Lysines. **(d)** Models were binned according to their agreement with XL-MS data from the standard search (well satisfied: 80-100 %, moderately satisfied: 20-80 %, poorly satisfied 0-20 %). Inter-link distance comparing both strategies and model subsets is depicted. A pLDDT cut-off of 50 was applied for both cross-linked Lysines. **(e)** Example of AF2-predicted SMC2-SMC4 heterodimer with Inter-links in standard and additional inter-links upon subgrouping. **(f)** Experimental cryoEM model of the proteasome and matching inter-links in the standard approach (ungrouped) and additional links obtained from inter-dependent grouping. **(g)** Histogram of distances obtained from (f). An inter-link level FDR of 0.01 was set for both standard and inter-dependent approaches.

