## Supplementary Note for "Enhancing Inter-link Coverage in Cross-Linking Mass Spectrometry through Context-Sensitive Subgrouping and Decoy Fusion"

**Supplementary Note: Generation of ground-truth dataset**

This ground-truth dataset was recently generated by our group to assist the development of an AI-based cross-link search engine Scout. Since this manuscript is still under preparation, we hereby supplement the experimental procedure and made the dataset available to the reviewers on ProteomeXchange under PXD042173 (, password: XXXXXX).

**Experimental procedure**

192 his-tagged human proteins were purified from E. coli using immobilized metal affinity chromatography and lyophilized. Reconstituted proteins were mixed in pairs following a defined mixing scheme. After incubation at 50 °C for 20 min, protein pairs were cross-linked at 0.1 - 1.5 mg/mL protein concentration with 0.2 - 1 mM DSSO for 30 min at RT. The reaction was quenched with 20 mM Tris-HCl pH 8.0. All pairs were mixed in individual groups of eight proteins, and proteins were denatured with 8 M Urea, alkylated, reduced, and digested into peptides using sequential LysC and over-night Trypsin digestion. Desalted peptides were separated offline using strong cation exchange chromatography, and selected fractions were subjected to 180 min LC-MS/MS measurements. RAW files were searched as described above with DSSO cross-linker specificity using a database containing all 192 mixed, 110 additional entrapment, and 302 reversed decoy proteins plus potential contaminants (targets and reversed decoys). All cross-links matching contaminant entries were removed prior to data analysis. True positives and false positives were assigned at the peptide level to account for the partial homology of proteins in different groups.

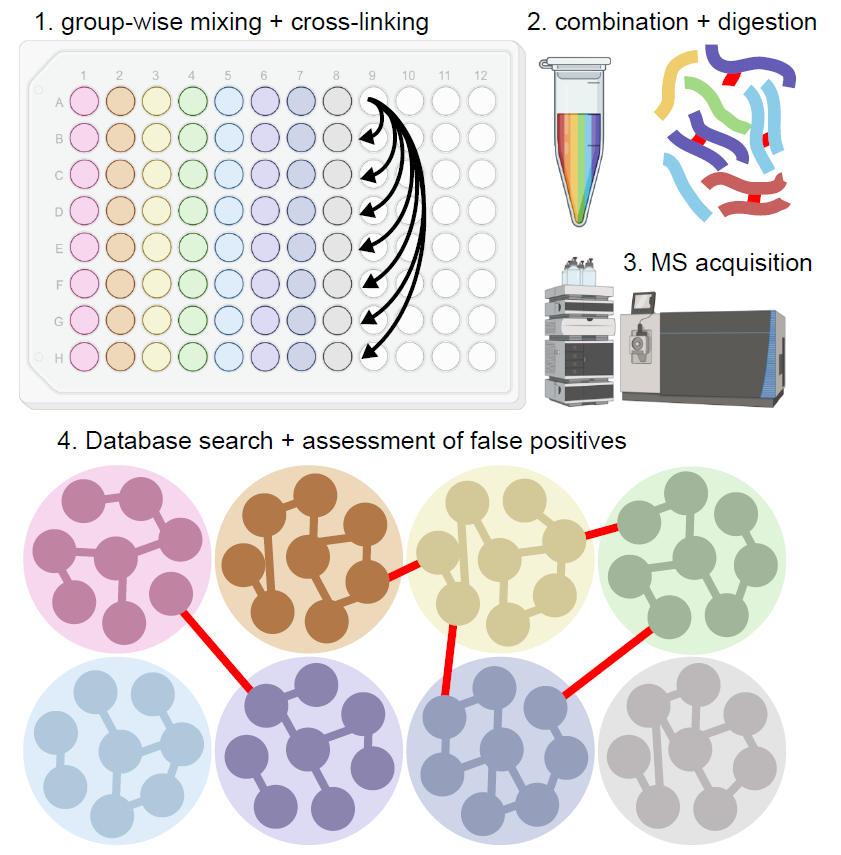

***Figure: A ground-truth dataset to evaluate context-sensitive subgrouping.* (a)** Experimental design of ground-truth dataset. Proteins were mixed in pairs, heat-treated to induce interactions and cross-linked with DSSO. This yields ground-truth true (edges within group) and false positives (edges between groups, highlighted red) according to the pre-defined mixing scheme.
